# TRIBES: A user-friendly pipeline for relatedness detection and disease gene discovery

**DOI:** 10.1101/686253

**Authors:** Natalie A. Twine, Piotr Szul, Lyndal Henden, Emily P. McCann, Ian P. Blair, Kelly L. Williams, Denis C. Bauer

## Abstract

**Summary:** TRIBES is a user-friendly pipeline for relatedness detection in genomic data. TRIBES is the first tool which is both accurate up to 7^th^ degree relatives (e.g. third cousins) and combines essential data processing steps into a single user-friendly pipeline. Furthermore, using a proof-of-principle cohort comprising amyotrophic lateral sclerosis cases with known relationship structures and a known causal mutation in *SOD1*, we demonstrated that TRIBES can successfully uncover disease susceptibility loci. TRIBES has multiple applications in addition to disease gene mapping, including sample quality control in genome wide association studies and avoiding consanguineous unions in family planning.

**Availability:** TRIBES is freely available on GitHub: https://github.com/aehrc/TRIBES/

**Supplementary information:** XXXX

## 4. Introduction

Accurately classifying the degree of relatedness between pairs of individuals has important applications, including empowering disease gene discovery in linkage analyses (Teare et al. 2005), removal of relatives in genome wide association studies (GWAS) (Voight & Pritchard 2005) and avoiding consanguineous unions which lead to poor health outcomes (i.e. family planning) (Shalev 2019). Several tools exist to infer the degree of relatedness between individuals using two main approaches: genomic regions that are identical-by-descent (IBD) and population allele frequency-based tools. GERMLINE (Gusev et al. 2009) and Refined IBD (Browning & Browning 2013) identify genomic segments that have been inherited from a common ancestor IBD, which can be parsed to additional tools such as ERSA (Li, Glusman, Hu, et al. 2014) to infer relationships. On the other hand, KING (Manichaikul et al. 2010) and PLINK (Purcell et al. 2007) use population allele frequencies to infer relatedness measures.

A key benefit of allele frequency-based tools is the fast compute time and minimal data pre-processing, however these tools show poor accuracy in identifying relatives more distant than 3rd degree. In contrast, IBD segment-based tools demonstrate significantly better accuracy for distant relatedness (>3rd degree), although often require the use of additional tools to perform ad-hoc data pruning, phasing of chromosomes and IBD inference.

Currently no tools are available that are reliable beyond 3rd degree and combine the necessary data processing steps for accuracy and ease of use. Being able to accurately and routinely perform relationship classification is especially relevant with recent increases in large-cohort whole genome sequencing (WGS) studies, which in turn increases the risk of related samples within a study. Accurate relationship classification is also imperative for linkage analysis to identify disease linked loci.

To address this we have developed a user-friendly pipeline, TRIBES, which combines data pruning, phasing of genomes, IBD segment recovery, masking of artefactual IBD segments and accurate relationship estimation into one tool. We demonstrate the accuracy of TRIBES on a simulated population, consisting of 1,290 samples and 18,480 known relationship pairs, as well as the tool’s ability to create novel insights using an amyotrophic lateral sclerosis (ALS) data set (Henden et al. 2019).

## 4. Methods

The input data to TRIBES is a quality control filtered, joint sample variant call format (VCF) file. TRIBES then follows the steps below for relatedness inference, where the full pipeline is described in detail in the Supplementary Material.

3. Variants in the VCF file are filtered according to quality control metrics using bcftools (v1.9) (Li et al. 2009; Li 2011).
3. The filtered VCF is then phased using BEAGLE (v4.1) (Browning & Browning 2007) with or without a reference dataset.
3. The phased VCF is converted to PLINK map format with vcftools (v0.1.16) from which IBD segments are then estimated using GERMLINE (v1.5.3) (Gusev et al. 2009).
3. IBD segments located within regions with high amounts of artefactual IBD are masked and segment endpoints are adjusted.
3. Adjusted IBD segments are then summed to estimate relationships between pairs of individuals.
3. TRIBES returns the relatedness estimates for all pairs of individuals as well as result files for all intermediate analysis steps.

## 3. Results

### 3.1 TRIBES is more accurate than KING given a fixed workflow

Haplotype data was generated for 18,480 pairs of related individuals from a simulated 15-generation pedigree and used to compare the accuracy of TRIBES (v 1.0.0) and KING (v. 2.0.0) (Figure 1, Supplementary Material). Given a fixed workflow we demonstrated that TRIBES is more accurate than KING at 3rd through to 15th degree relatives. Particularly beyond 4th degree relatives, the accuracy of KING drops off precipitously, calling 41.27% of known 5th degree and 10.17% of 6th degree relationships correctly, while TRIBES called 56.06% and 43.64% of 5th and 6th degree relationships correctly. We have also applied TRIBES to an ALS disease cohort comprising known relationships (Henden et al. 2019) where we demonstrated 99% accuracy (allowing 1 degree of error) up to 7th degree relatives, where 5th degree relatives showed 73.02% accuracy and 6th degree relatives showed 61.02% accuracy.

**Figure 1.**
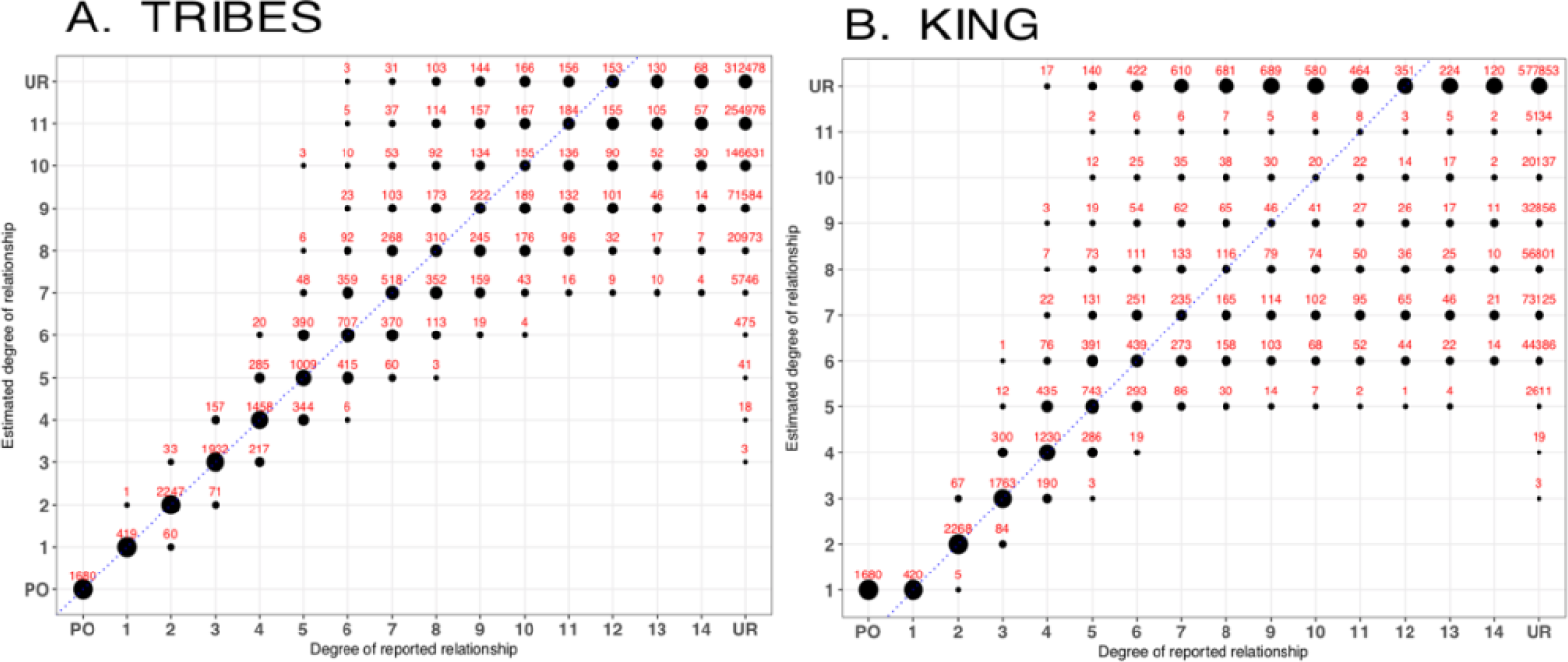
Performance of relationship estimation in 30 simulated pedigrees with (A) TRIBES 1.0.0 and (B) KING 2.0.0. The size of the circles indicates the percentage of pairs whose estimated degrees of relationship are identical to the reported relationship. Absolute count of relationships is shown in red above the circle. The dashed blue line along the diagonal indicate relationship pairs for which the reported and estimated degree are the same. PO: parent offspring. UR: unrelated individuals.

Furthermore, TRIBES discovered novel relationships in the ALS disease cohort (Henden et al. 2019).

### 3.2 TRIBES is user-friendly due to built-in data pruning and flexible workflow

TRIBES includes built-in data pruning and phasing that would otherwise need to be performed using multiple tools prior to IBD analysis with GERMLINE and relationship estimation. The entire analysis is performed using a single command that includes an option for multithreading to reduce computation time. TRIBES leverages workflow tool Snakemake (Köster & Rahmann 2012), which provides for reproducibility and seamless scalability to server, cluster, grid and cloud environments, as well as allows for flexible customisation of the data processing pipeline steps and their parameters.

### 3.3 Masking of artefactual IBD enables disease locus discovery

We observed regions of the genome which had disproportionately high amounts of shared IBD (Supplementary Figure 2A and B), in accordance with previous studies (Li, Glusman, Huff, et al. 2014). These regions were generally consistent between the unrelated 1000 Genomes ‘EUR’ cohort (Supplementary Figure 2A) and an independent ALS pedigree (Henden et al. 2019) (Supplementary Figure 2B), indicating these regions are indeed artefactual. Hence, TRIBES incorporates masking of artefactual IBD prior to relationship estimation, similarly to ERSA 2.0 (Li, Glusman, Huff, et al. 2014). This not only significantly improves the accuracy of relationship estimates, but also allows for disease gene mapping, where a locus with high amounts of IBD sharing is observed over the known ALS gene *SOD1* (Supplementary Figure 2c), where all individuals in this cohort carry one of three disease-causing *SOD1* mutations (Henden et al. 2019).

## 4. Conclusion

We have developed a novel relatedness tool, TRIBES, which addresses the need for a relatedness discovery platform that is accurate beyond 3^rd^ degree relatives, easy-to-use and flexible. TRIBES is an end-to-end platform that can process large cohorts of WGS data, incorporating multiple tools and methodologies into a single, straightforward pipeline. Furthermore, TRIBES can substantially narrow the search space for disease loci in cohorts of related samples, making it an essential tool for researchers investigating the genetic origins of disease. TRIBES has further utility in both sample quality control prior to GWAS and for avoiding consanguineous unions in family planning.

## Acknowledgements

We thank Carolyn Cecere and Lorel Adams for their assistance in compiling *SOD1* family information and Elisa Cachia and Dr Sarah Furlong for providing patient materials, clinical and technical assistance.

