## Supplementary material for "TRIBES: A user-friendly pipeline for relatedness detection and disease gene discovery"

### TRIBES supplementary material:

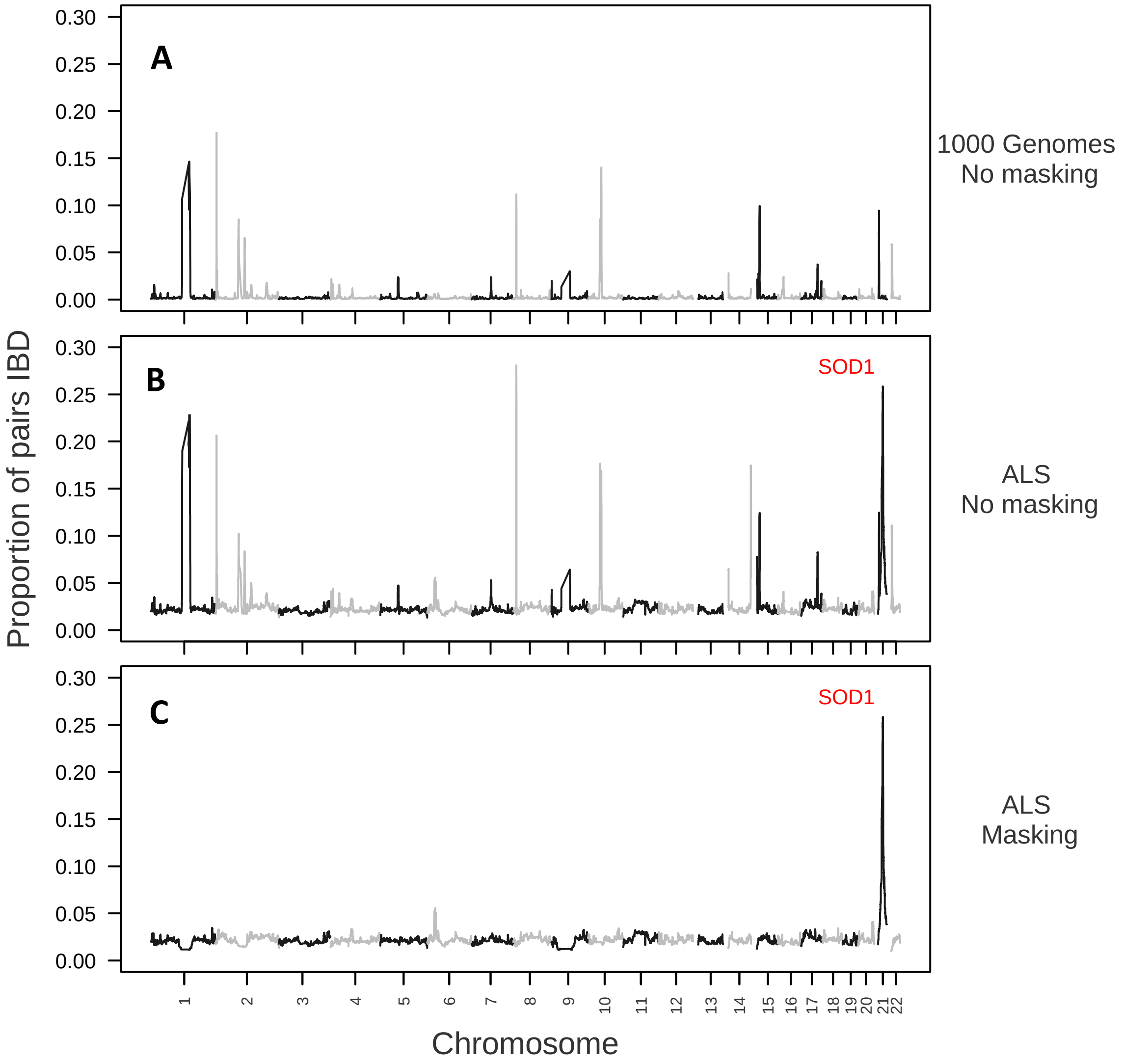


**Supplementary Figure 2.**

Genome-wide distribution of the proportion of pairs IBD before (A, B) and after (C) masking with TRIBES. Figure (A) shows the proportion of pairs IBD for the 1000 Genomes unrelated 'EUR' cohort, while (B) and (C) show the proportions for an independent ALS cohort (n=88) (Henden et al. 2019). All samples in the ALS cohort have a mutation in the known ALS gene, *SOD1*. The *SOD1* locus is located on chromosome 21 and highlighted by a red line in Figure 2B and 2C.

**Steps involved in TRIBES:**

1. ***Filtering on quality metrics within VCF file***

   An input VCF file is filtered to include only biallelic variants which pass certain quality control metrics using bcftools (v1.9) (Li et al. 2009; Li 2011). Each variant needs to pass all of the following criteria: MQ>59, MQRankSum >-2, DP>20, DP<100, QD>15, BaseQRankSum>-2, SOR<1. Variants which do not have a 'PASS' in the FILTER field are also excluded. Following this, variants with a minor allele frequency (MAF) < 1% as well as variants in linkage disequilibrium (LD) ($R_2 > 0.95$) are filtered out. MAF and LD values are calculated using the phase 3 1000 Genomes EUR (European) cohort (including only unrelated 1000 Genomes samples) (Auton et al. 2015).

2. ***Phasing***

   The filtered VCF is then phased using BEAGLE (v4.1) (Browning & Browning 2007) with the 1000 Genomes 'EUR' data as a reference dataset (Auton et al. 2015), where missing variants are imputed. After phasing, vcftools (v0.1.16) (Danecek et al. 2011) is used to create PLINK .ped and .map files, including a custom script to interpolate genetic distance based on a genetic map from (Auton et al. 2015).

3. ***IBD segment calculation***

   GERMLINE (v.1.5.3) (Gusev et al. 2009) is then run on the resultant files, with the options '-bits 128 -min_m 1.5 -err_het 1 -err_hom 2 -g_extend –w_extend' to identify IBD segments shared between pairs of individuals.

4. ***Masking for artefactual IBD***

   In order to account for artefactual IBD present in datasets, we built a custom masking function to post-process IBD segments recovered by GERMLINE in step 3. This function adjusts the endpoints of IBD segments if those segments overlap with loci that have high amounts of IBD sharing in the unrelated 1000 Genomes 'EUR' reference population, reflecting artifactual IBD (Supplementary Table 1, Supplementary Figure 1a and 1b). This function utilises the methodology outlined in the 'Genomic Region Masking' section of ERSA 2.0 publication (Li et al. 2014).

| chromosome | begin_position | end_position |
|---|---|---|
| 1 | 11,976,168 | 14,507,057 |
| 1 | 107,871,447 | 160,529,058 |
| 2 | 1,247,922 | 6,916,359 |
| 2 | 78,301,690 | 118,902,913 |
| 2 | 134,582,082 | 138,119,920 |
| 2 | 191,886,543 | 199,830,770 |
| 4 | 3,685,427 | 4,032,938 |
| 4 | 7,148,852 | 12,391,898 |
| 4 | 29,334,842 | 37,555,938 |
| 5 | 63,877,512 | 74,977,117 |
| 7 | 71,388,036 | 77,594,411 |
| 8 | 8,073,336 | 16,397,104 |
| 9 | 1,161,645 | 2,507,038 |
| 9 | 22,844,440 | 91,209,596 |
| 10 | 30,516,851 | 61,488,215 |
| 14 | 19,006,459 | 24,049,962 |
| 14 | 106,043,805 | 107,107,217 |
| 15 | 26,129,659 | 35,040,766 |
| 16 | 17,660,644 | 24,384,636 |
| 17 | 55,274,367 | 66,318,466 |
| 17 | 77,350,369 | 77,911,445 |
| 21 | 15,577,434 | 20,832,668 |
| 22 | 16,554,781 | 23,222,208 |

**Supplementary Table 1.** Regions of genome from unrelated 1000 Genomes 'EUR' population that have high amounts of IBD sharing, reflecting artifactual IBD. These regions are used as a reference set of regions to mask in step 4 of TRIBES pipeline,

5. ***Relationship estimation***

The lengths of IBD segments shared between pairs are then summed to calculate the proportion of the genome with zero alleles inferred IBD (IBD0) for each pair. This enables TRIBES to estimate degrees of relationship, according to expected IBD0 ranges for each pair, detailed in Supplemental Table 2. Expected IBD0 values are taken from previously published work (Ramstetter et al. 2017).

6. ***TRIBES returns result files***

After successful completion of TRIBES, a number of intermediate and result files are returned. Estimated relatedness for every sample pair is returned as .csv file in the output directory. If a file containing true (reported) degrees was supplied to TRIBES, an .html file is returned which displays the accuracy of the estimated degree relative to the reported degree.

| **Degree** | **IBD0 values that map to Degree** | **Relationship** |
|---|---|---|
| 0 | <0.1 | Monozygotic (MZ) twin |
| 1 | <0.1<br>[0.1 - 0.366] | Parent-child<br>Full sibling (excluding MZ twin) |
| 2 | [0.366 – 0.646] | Grandparent – grandchild<br>Avuncular<br>Double-cousin<br>Half-sibling |
| 3 | [0.646 – 0.823] | First cousin<br>Great-grandparent<br>Grand-avuncular<br>Half-avuncular |
| 4 | [0.823 – 0.912] | First cousin once removed<br>Great-great-grandparent<br>Great-grand-avuncular<br>Half-grand-avuncular |
| 5 | [0.912 - 0.956] | First cousin twice removed<br>Second cousin<br>GGG-grandparent |
| 6 | [0.956 – 0.978] | Second cousin once removed |
| 7 | [0.987 – 0.989] | Second cousin twice removed<br>Third cousin |
| 8 | [0.989 – 0.994] | Third cousin once removed |

**Supplementary Table 2.** Degrees of relatedness with the ranges of genome proportions inferred to be IBD0 for each degree. Example relationships are listed for each degree, which is non-exhaustive for each degree. This table is an abridged version of that published in (Ramstetter et al. 2017).

**Generation of simulated pedigree**

We simulated 15 generation pedigrees of WGS data, using founders from unrelated 1000 Genomes samples of European (EUR) descent (Auton et al. 2015). In total we simulated 30 pedigrees, each containing 18,480 related pairs. Genotypes of non-founders were obtained by simulating meiosis, where recombination was introduced according to a recombination map (McVean et al. 2004). Recombination was modelled using an exponential distribution with mean equal to 1 Morgan. We added de-novo mutations at a rate of 1.1e-8 per base per generation. The simulated pedigree dataset is available for download: https://csiro-tribes.s3-ap-southeast-2.amazonaws.com/downloads/examples/TFEur.tar.gz

**TRIBES analysis on the simulated data**

- **Step 1.** SNPs with MAF < 1% and SNPs in high LD (R2 > 0.95) were removed. Filtering on other quality control metrics was not performed as this information was not generated as part of the simulated VCF file.
- **Step 2.** The simulated haplotype data was then phased using BEAGLE.
- **Steps 3-6.** Analysis was performed for these steps according to the default pipeline parameters.

**KING analysis on the simulated data**

No filtering was performed on the simulated haplotype data prior to analysis with KING(v.2.0.0) (Manichaikul et al. 2010). However, PLINK (v.1.90) (Purcell et al. 2007) was used to reformat the simulated data for input into KING. The following command lines were implemented.

*plink –vcf fakefamily.vcf.gz --allow-extra-chr --make-bed --out FakeFam*
*king -b FakeFam.bed --kinship --prefix FakeFam*

**Amyotrophic lateral sclerosis (ALS) cohort**

The IBD analysis of the ALS cohort used in this study has been previously described in (Henden et al. 2019) while the processing of whole genome sequencing data for this cohort is described in (Henden et al. 2019).

**Snakemake and R package**

The full pipeline is run using Snakemake (v. 5.4.5) (Köster & Rahmann 2012) to enable reproducibility as well R (Version 3.5.1) (Rstudio Team 2015) and Python (3.7). Users can adjust the filtering/processing parameters within the Snakemake 'config.yaml' file for any step in TRIBES.

**Compute resources used in analysis**

Analysis of the simulated data with TRIBES and KING was performed on a High Performance Computing cluster, which consists of 230 Dell PowerEdge M630 servers with 128 GB of memory and dual 10 core Intel Xeon E5-2660 V3 CPUs. For the purpose of TRIBES analysis 22 CPU cores were utilized concurrently (one core per chromosome) for all steps except for phasing which used a full node (20 CPU cores) per chromosome.
Detailed information on how to run TRIBES on an HPC cluster or local machine can be found here https://github.com/aehrc/TRIBES.

Rstudio Team, 2015. RStudio: Integrated Development for R. *[Online] RStudio, Inc., Boston, MA.*, p.RStudio, Inc., Boston, MA.
